# The yeast DENN domain protein Avl9 contributes to recycling and sorting of endosomal cargos

**DOI:** 10.64898/2026.02.08.704655

**Authors:** Daniel J. Rioux, Samreen Manj, Derek Prosser

## Abstract

In yeast and humans, the conserved DENN-domain (<u>D</u>ifferentially <u>E</u>xpressed in <u>N</u>ormal and <u>N</u>eoplastic tissue) protein Avl9 is thought to play roles in membrane traffic and secretion, but its precise function remains poorly defined. Since DENN-containing proteins are associated with Rab GTPase function, we sought to understand Avl9 function in the context of Rab regulation. Here, we show that Avl9 localizes to peripheral punctae that are consistent with secretory vesicles. Moreover, we demonstrate genetic interactions and co-localization between Avl9 and numerous Rabs in the secretory and endosomal pathways, suggesting a potential function at the interface of secretion and recycling. Consistent with this role, *avl9*Δ results in defective recycling of the endosomal cargo Snc1 but does not alter plasma membrane delivery of an endocytosis-defective Snc1^EN-^ mutant, suggesting that Avl9 is not directly involved in secretory traffic from the TGN to the plasma membrane. The *avl9*Δ recycling defect is exacerbated by the additional loss of *RCY1* or *SNX4*, but not *VPS35*. Each of these three genes contributes to a distinct endosomal recycling pathway, indicating that Avl9 acts in conjunction with multiple recycling pathways.

**Summary Statement:** In this study, Rioux *et al*. describe a role for the DENN domain protein Avl9, previously thought to regulate secretion, as a novel factor involved in recycling of cargos from endosomal compartments.

## Introduction

Vesicular trafficking is a process critical to maintaining the composition, identity, and function of organelles throughout all eukaryotic cells, including the budding yeast *Saccharomyces cerevisiae* (Zerial and McBride, 2001). Distinct trafficking pathways are defined by the origin and destination of the vesicle, and involve unique sets of proteins for cargo selection and vesicle biogenesis. The secretory pathway routes newly synthesized proteins from the endoplasmic reticulum through the Golgi to the plasma membrane (PM) or, alternatively, to the endosome. Vesicles containing cargos from the PM are internalized via clathrin-mediated and clathrin-independent endocytic pathways, permitting cargo delivery to endosomes and the *trans*-Golgi network (TGN) (Kaksonen and Roux, 2018; Rioux and Prosser, 2023). At endosomes, cargos enter the endolysosomal pathway to be directed through a series of maturing compartments including early endosomes (EEs), late endosomes (LEs), and multivesicular bodies (MVBs) before delivery to the vacuole/lysosome (Babst et al., 1997; Henne et al., 2011; Hickey and Wickner, 2010; Raymond et al., 1992; Vida and Emr, 1995). At each stage of the endolysosomal pathway, cargo can be directed back to the PM and Golgi through a series of recycling pathways (Best et al., 2020; Hettema et al., 2003; Laidlaw et al., 2022; Ma and Burd, 2020; MacDonald and Piper, 2017). Collectively, these pathways form a complex network, requiring many distinct proteins to properly sort cargos, form vesicles, and ensure accurate targeting to the destination compartment.

Small GTPases of the Rho, Arf and Rab families play key roles in regulating vesicle formation, targeting and fusion (D’Souza-Schorey and Chavrier, 2006; Gillingham and Munro, 2007; Jaffe and Hall, 2005). Small GTPases cycle between active, GTP-bound and inactive, GDP-bound states, where the active state enables association with effector proteins that carry out downstream functions (Bos et al., 2007). This cycling process, along with their recruitment to and regulation at distinct compartments, is regulated by two classes of protein: guanine nucleotide exchange factors (GEFs) promote the release of GDP from an inactive GTPase, leading to GTP binding and conformational changes to achieve an active state (Müller and Goody, 2017). Conversely, GTPase-activating proteins (GAPs) bind to active GTPases to facilitate hydrolysis of GTP to GDP, leading to inactivation.

Rab GTPases are involved in all stages of membrane traffic, and are highly conserved through eukaryotic evolution. Rabs perform broad roles within the cell through their interactions with distinct recruitment factors and effectors. Recruitment of Rab effectors to target compartments plays a valuable role in establishing and maintaining organelle identity (Cai et al., 2007). Effector interactions allow Rabs to direct targeting of vesicles to the correct destination through engagement with myosin motor proteins that move vesicles along the actin cytoskeleton and with tethering complexes found on specific organelles (Hickey and Wickner, 2010; Mima, 2018; Ostrowicz et al., 2010). Additionally, Rab interactions with SNARE proteins are critical to driving fusion of vesicles with their target membranes (Cai et al., 2007; Grote and Novick, 1999).

A family of proteins characterized by a DENN (differentially expressed in normal and neoplastic cells) domain contains members that function either as GEFs or GAPs toward small GTPases, especially (but not exclusively) within the Rab family (Jansen and Hurley, 2023; Marat et al., 2011). Budding yeast contains several DENN domain proteins, including the evolutionarily conserved Avl9. *AVL9* was first described in yeast as a gene that showed synthetic lethality in an *apl2*Δ *vps1*Δ background defective in Golgi sorting and exit (Harsay and Schekman, 2007). Avl9 depletion in *apl2*Δ *vps1*Δ resulted in defective secretion of numerous cargos. Further characterization of Avl9 function has remained difficult, and *avl9*Δ does not have any obvious phenotypes reported in the literature. Here, we demonstrate that Avl9 localizes to secretory vesicle-like structures and that Avl9 co-localizes with numerous Rabs. Further, we show that *avl9*Δ results in mislocalization of the v-SNARE Snc1, which typically undergoes rounds of recycling between the endosomal system and the PM but is instead retained on internal compartments in the absence of Avl9. We also demonstrate that disruption of Snc1 recycling in *avl9*Δ is exacerbated by the loss *RCY1* or *SNX4.* Taken together, our data suggest that Avl9 contributes to sorting and secretion of endosomal cargos through multiple recycling pathways.

## Results

### Avl9 localizes to secretory vesicle-like structures

To gain insight into possible functions of Avl9, we initially examined the subcellular localization of Avl9-GFP, expressed from its endogenous promoter, by fluorescence microscopy (Fig 1A). We found that Avl9-GFP showed diffuse cytosolic signal and additionally localized to bright punctae with distinct cell cycle-dependent distribution. In unbudded cells (G1 phase), Avl9-GFP punctae localized primarily to the cell periphery, with a small number of internal structures. In small-budded cells (S phase), we observed Avl9-GFP punctae that polarized towards the nascent bud site; these punctae strongly polarized to the daughter cell in medium-budded cells (G2 phase). Finally, Avl9-GFP localized almost exclusively to the bud neck, with very few punctae in large-budded cells (M phase). Overall, this localization pattern is consistent with that of other late-secretory trafficking proteins such as Sec4 and strengthens earlier findings suggesting a role for Avl9 in secretion (Boyd et al., 2004; Goud et al., 1988; Harsay and Schekman, 2007).

**Figure 1:**
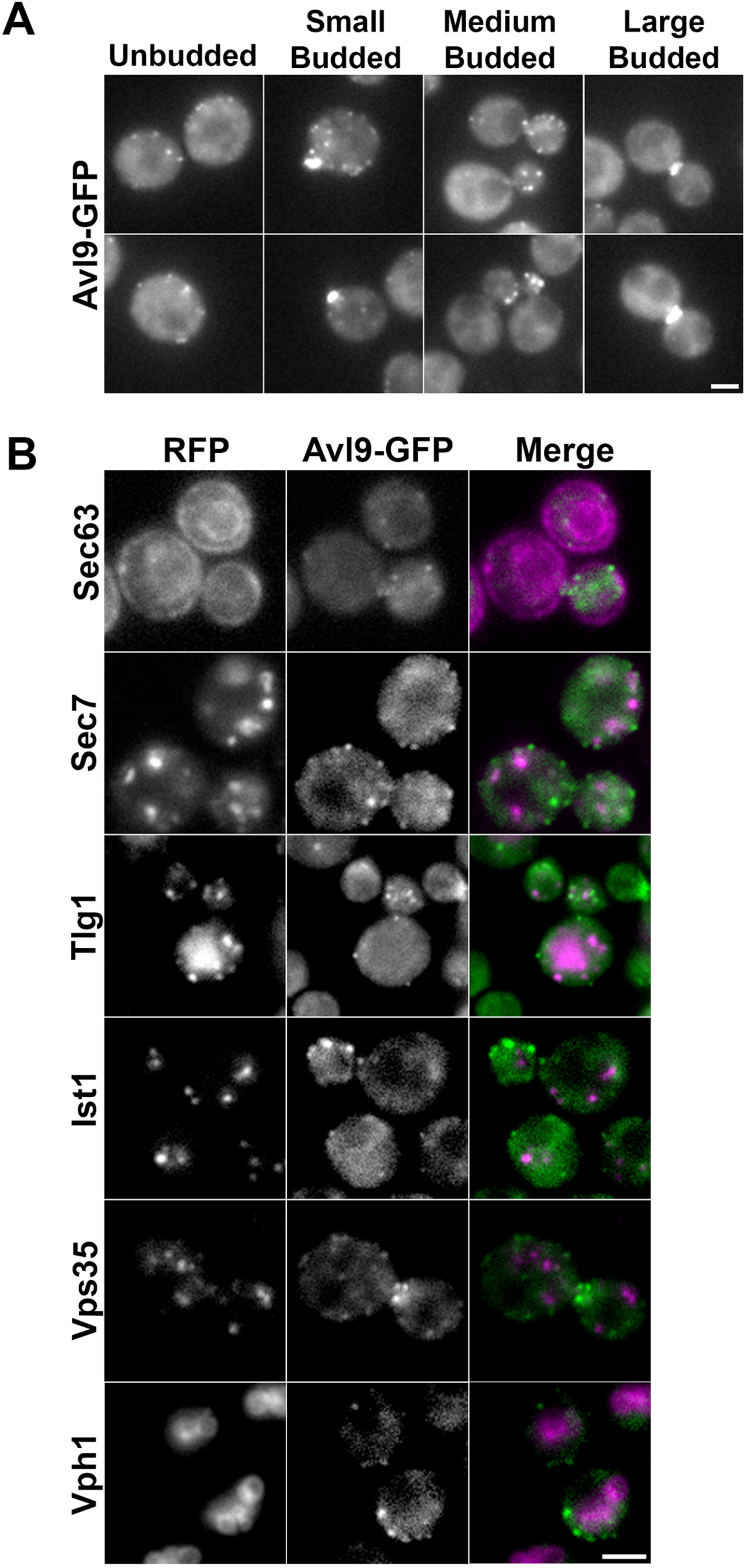
Subcellular localization of Avl9-GFP. (A) Cells expressing Avl9-GFP were imaged by fluorescence microscopy. Images show endogenously-tagged Avl9-GFP localization in unbudded as well as small-, medium-, and large-budded cells. (B) Two-color fluorescence microscopy of cells expressing Avl9-GFP an organelle marker endogenously tagged with mScarlet1 (Sec63, Sec7, Ist1, Vps35, Vph1) or with mCherry and expressed from a plasmid (Tlg1). Scale bar = 2 μm.

Next, we sought to determine whether Avl9-GFP localized to any major organelles by generating Avl9-GFP strains with mScarlet1-tagged organelle marker proteins expressed from their endogenous loci. These markers were chosen from a previously validated set of proteins residing at the ER (Sec63), TGN (Sec7), late endosomes (Ist1), multivesicular bodies (Vps35) and vacuoles (Vph1) (Zhu et al., 2019). Additionally, we transformed Avl9-GFP cells with a plasmid expressing the t-SNARE mCherry-Tlg1 as an early endosomal marker (Vida and Emr, 1995; Wiederkehr et al., 2000). We then imaged these strains using simultaneous two-color fluorescence microscopy, but failed to observe significant co-localization of Avl9-GFP with any of the organelle markers examined, regardless of the stage in the cell cycle (Fig. 1B). While the lack of co-localization with Sec63, Ist1, Vps35, or Vph1 was unsurprising given the putative role for Avl9 in late stages of secretion, we also observed no co-localization between Avl9 the TGN marker Sec7, which was unexpected because the TGN is the site from which secretory vesicles originate. These data suggest that Avl9-positive punctae may be vesicular in nature and that Avl9 likely arrives on secretory vesicles after scission and departure from their point of origin.

### Avl9 co-localizes and shows genetic interactions with numerous Rabs

We next sought to assess localization of Avl9 to vesicles along all major trafficking pathways within the cell. We chose the Rab family of small GTPases as markers for the pathways for several reasons. First, Rab GTPases are well-studied proteins that act on vesicles operating in distinct pathways to confer directionality to membrane traffic (Hickey and Wickner, 2010; Mima, 2018; Ostrowicz et al., 2010). This allows us to have a singular class of proteins to make localization comparisons against. Second, the major structural feature of Avl9 is a DENN domain (Harsay and Schekman, 2007). DENN domain proteins are a broad class of conserved proteins that interact with several small GTPase families, including Rabs, as effectors or regulators (Marat et al., 2011). We reasoned that if Avl9 is a DENN domain protein involved in trafficking, it may be recruited to Rab-positive vesicles. We generated a genomically-encoded Avl9-mCherry strain, transformed with plasmids carrying N-terminally tagged GFP-Rabs expressed from their endogenous promoters, and imaged cells using simultaneous two-color fluorescence microscopy. Surprisingly, we found that Avl9-mCherry co-localized with numerous Rabs (Fig. 2A), both expected and unexpected based on our assessment of Avl9 coincidence with organelle markers (Fig. 1B). Given the localization pattern of Avl9-GFP, we anticipated the observed co-localization with Sec4; however, we also observed coincidence with the early secretory Rabs Ypt1 and Ypt31, and early endosomal Rab5 isoforms Ypt52 and Ypt10 (Buvelot Frei et al., 2006; Jedd et al., 1995; Singer-Krüger et al., 1994). Co-localization with Ypt52 and Ypt10 was notable because Avl9 showed little to no coincidence with the remaining two Rab5 isoforms, Vps21 and Ypt53 (Buvelot Frei et al., 2006; Singer-Krüger et al., 1994).

**Figure 2:**
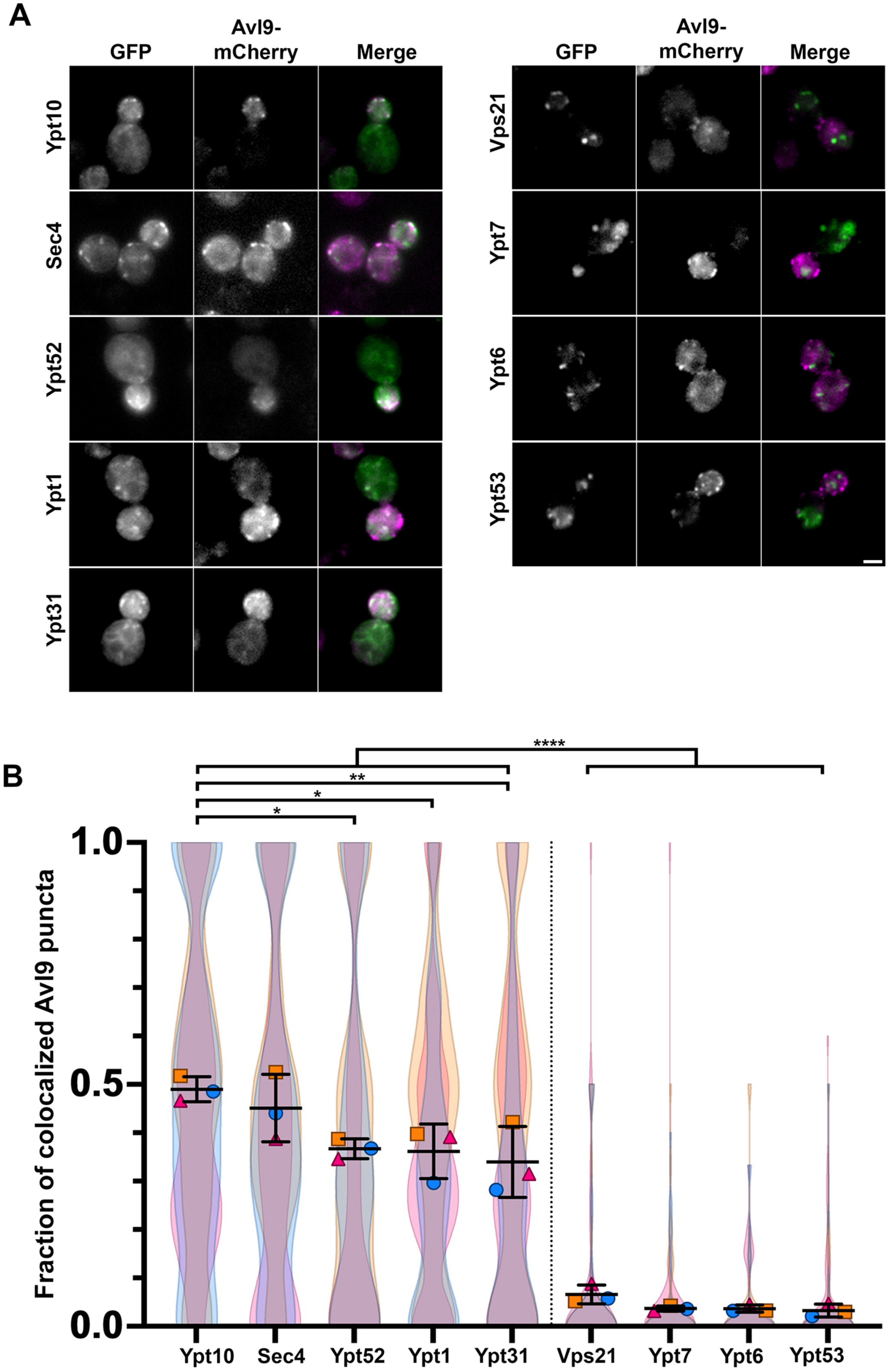
Colocalization of Avl9 with Rabs. (A) Two-color fluorescence microscopy of cells expressing endogenously-tagged Avl9-mCherry and plasmid-borne GFP-tagged Rabs. Scale bar = 2 μm. (B) SuperPlots showing quantification of trial-level means ± SD overlaid on violin plots (color matched to trial-level summary) of each trial for the fraction of Avl9-mCherry punctae that co-localize with GFP-Rabs. Rabs are ordered based on the highest fraction of Avl9 co-localization. One-way ANOVA with Tukey’s multiple comparisons test. n=3 biological replicates, with 35-49 cells measured per condition for individual trials (* p<0.05, ** p<0.01, **** p<0.0001).

To better understand the co-localization between Avl9 and Rabs, we sought to quantify our findings. While many quantitative co-localization analyses rely on Pearson or Spearman correlations, or on Mander’s coefficients, we were concerned that these methods may leave out valuable information or distinctions given the dynamic nature of vesicle trafficking and the cytoplasmic and membrane-associated distribution of Rabs and Avl9 (Adler and Parmryd, 2010; Jimenez et al., 2022). Instead, we manually counted all Avl9-mCherry punctae in each cell and determined the fraction of total Avl9-containing structures that coincided with a GFP-Rab puncta. We then ordered the Rabs based on how well Avl9 co-localized with them (Fig. 2B). The Rabs showing the strongest co-localization fraction with Avl9 were Ypt10 (0.48 ± 0.026) and Sec4 (0.45 ± 0.069), followed by Ypt52 (0.36 ± 0.021), Ypt1 (0.36 ± 0.057) and Ypt31 (0.33 ± 0.073). The remaining Rabs Vps21 (0.06 ± 0.019), Ypt7 (0.03 ± 0.01), Ypt6 (0.03 ± 0.01) and Ypt53 (0.03 ± 0.01) showed substantially lower co-localization fractions, suggesting poor coincidence with Avl9.

Our finding that Avl9 broadly co-localizes with Rabs in both the secretory pathway and the early stages of the endolysosomal pathway suggest that the role of Avl9 in membrane traffic may not be restricted to TGN-to-PM secretory transport. To better understand the relationship between Avl9 and Rabs, we attempted to identify genetic interactions arising from combined *AVL9*/*RAB* loss-of-function mutations. We first deleted *AVL9* in the temperature-sensitive *sec4-8* mutant given the critical role Sec4 plays in the late secretory pathway (Walworth et al., 1989). Plate-based growth of WT, *sec4-8* and *sec4-8 avl9*Δ showed that *avl9*Δ exacerbated the temperature sensitivity phenotype of the *sec4-8* mutant, resulting in weak growth even at normally permissive temperatures and demonstrating a relationship to secretion separate from the previously-identified interactions with *VPS1* and *APL2* (Fig. 3A) (Harsay and Schekman, 2007). We also utilized the temperature sensitive *bet3-1* allele to assess genetic interactions between *AVL9* and *BET3,* a core component of the TRAPP complexes which act as GEFs for Ypt1 and Ypt31/32, and saw no changes in temperature sensitivity (Supplemental Fig. S1A) (Lipatova and Segev, 2019; Menon et al., 2006; Sacher et al., 2001; Thomas and Fromme, 2016; Wang et al., 2000). We next investigated the relationship between the early endosomal yeast Rab5 isoforms and Avl9 by generating single, double and triple deletion mutants of *VPS21, YPT52,* and *YPT53* with or without *AVL9* (Singer-Krüger et al., 1994). We then grew all strains on plates with increasing concentrations of monensin, an ionophore that facilitates transport of Na^+^ ions across the membrane of the TGN and the early endosome, disrupting the pH and function of these compartments (Gustavsson et al., 2008). We reasoned that since *vps21*Δ displays monensin sensitivity, loss of *AVL9* might enhance this phenotype if it functions on endosomal vesicles as suggested by its co-localization with Ypt52 and Ypt10. Our growth assays confirmed this genetic interaction, with loss of *AVL9* increasing monensin sensitivity in *vps21*Δ, *vps21*Δ *ypt52*Δ, and *vps21*Δ *ypt52*Δ *ypt53*Δ (Fig. 3B). We similarly tested Ypt10, which is closely related to the Rab5 isoforms, but observed no phenotypes associated with *ypt10*Δ or genetic interactions with *avl9*Δ (Supplemental Fig. S1B).

**Figure 3:**
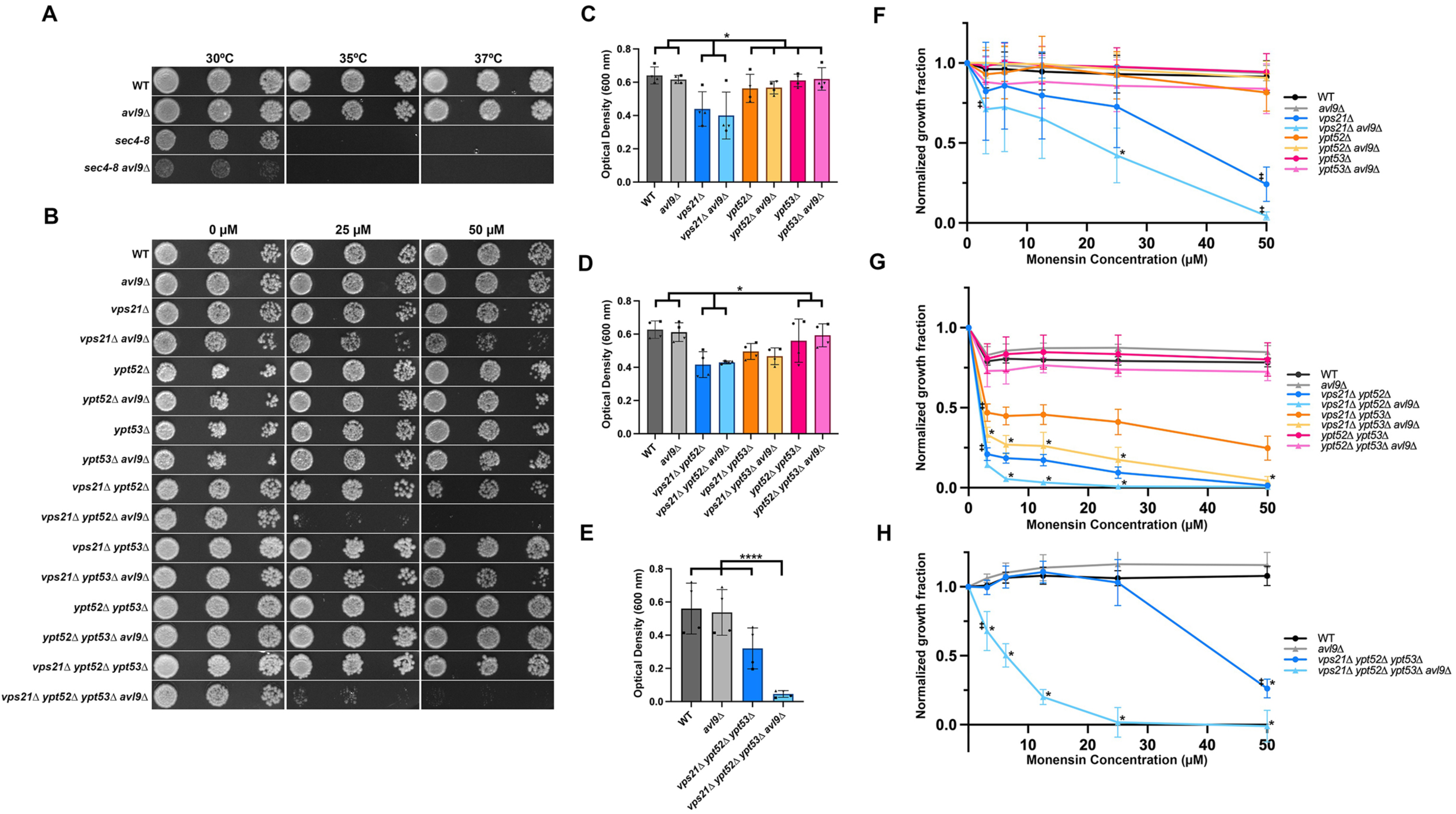
Genetic interactions between *AVL9* and Rabs. (A) WT and *sec4-8* cells with and without *AVL9* were grown to mid-logarithmic phase, serially diluted, and spotted onto YPD plates at the indicated temperatures for 3 d. (B) WT, *vps21*Δ, *ypt52*Δ*, ypt53*Δ, as well as Rab5 double and triple deletion mutants with and without *AVL9* were grown to mid-logarithmic phase, spotted onto YPD plates containing 0, 25, and 50 μM monensin, and imaged after 3 days growth at 30°C. (C-E) Endpoint growth assay of WT, *vps21*Δ, *ypt52*Δ*, ypt53*Δ, as well as Rab5 double and triple deletion mutants with and without *AVL9.* Stationary-phase cells were treated with 0, 3.125, 6.25, 12.5, 25, and 50 μM monensin in 96 well plates and grown for 18 h at 30°C. OD_600_ was then to assess culture density. Panels C-E show comparative growth of WT, *vps21*Δ, *ypt52*Δ*, ypt53*Δ, as well as Rab5 double and triple deletion mutants with and without *AVL9* at 0 μM monensin. Panels F-H show growth of WT, *vps21*Δ, *ypt52*Δ*, ypt53*Δ, as well as Rab5 double and triple deletion mutants with and without *AVL9* at each monensin concentration normalized to the control (0 μM) treatment for each strain. Two-way ANOVA with Tukey’s multiple comparisons test; (mean ± SD, n=3), ‡ represents the first point where RabΔ is significantly different from WT (p<0.05), while * represents points at which RabΔ is significantly different from RabΔ *avl9*Δ (p<0.05).

Given that some of the Avl9-dependent phenotypes were modest compared to controls, we sought to perform a quantitative assay to provide a more detailed understanding of this complex relationship. We used a 96-well plate-based endpoint assay to examine growth of monensin-treated Rab5Δ mutant strains with or without *avl9*Δ, which allowed us to measure growth by spectrophotometry (Hung et al., 2018). We analyzed growth after 18 h incubation at 30°C in two ways: first, we compared growth between strains at 0 µM monensin to assess baseline growth differences. Amongst the single Rab5 deletions, *vps21*Δ showed weaker growth that was not altered by the loss of *avl9*Δ, while neither *ypt52*Δ nor *ypt53*Δ showed any differences (Fig. 3C). In the double Rab5 deletion mutants, *vps21*Δ *ypt52*Δ showed weaker growth that was not altered by loss of *avl9*Δ, while the other mutants showed no significant growth defects (Fig. 3D). Surprisingly, *vps21*Δ *ypt52*Δ *ypt53*Δ showed no growth defect compared to WT, but the additional deletion of *AVL9* resulted in severely reduced growth (Fig. 3E). These findings are consistent with *VPS21* being the main Rab5 isoform in yeast Rab5, while *YPT52* and *YPT53* play more dispensable roles that may be able to compensate for lost functions caused by deletion of *AVL9*.

As a secondary analysis of the endpoint assays, we evaluated the effect of monensin on our Rab5 deletion mutants by normalizing the absorbance of each concentration to the baseline absorbance at 0 µM, thereby generating concentration-dependent growth curves. Analysis of monensin sensitivity revealed that among the Rab5 single delete mutants, *vps21*Δ and *vps21*Δ *avl9*Δ both showed a downward trend indicating sensitivity to monensin, with a significant difference beginning at 25 µM (Fig. 3F). The *vps21*Δ *ypt52*Δ and *vps21*Δ *ypt53*Δ mutants, with and without *AVL9,* all showed sensitivity to monensin at low concentrations, with *avl9*Δ increasing drug sensitivity (Fig. 3G). Finally, the *vps21*Δ *ypt52*Δ *ypt53*Δ mutant showed a surprising resistance to monensin, not becoming sensitive until the highest concentration while the *vps21*Δ *ypt52*Δ *ypt53*Δ *avl9*Δ mutants showed increased sensitivity at all concentrations (Fig. 3H). The loss of all three major Rab5 isoforms may lessen sensitivity to monensin because of pronounced perturbations to the endosomal system (Gustavsson et al., 2008; Singer-Krüger et al., 1994).

### Avl9 disrupts recycling and exocytosis of endosomal cargos

Given our results suggesting Avl9 involvement in trafficking within both secretory and endosomal pathways, we sought to determine whether loss of *AVL9* resulted in disruption of cargo transport. We chose yeast carboxypeptidase Y (CPY) because it is normally sorted from the TGN to the vacuole along the vacuole protein sorting (VPS) pathway, but can be missorted into secretory vesicles by deletion of *VPS* genes (Marcusson et al., 1994). One of these genes is *VPS4,* an ATPase that functions with the ESCRT-III complex to drive formation of multivesicular bodies (Babst et al., 1997; Harsay and Schekman, 2002). The CPY missorting phenotype allowed us to simultaneously assess disruption of cargos in both pathways using an assay in which WT, *avl9*Δ, *vps4*Δ, and *avl9*Δ *vps4*Δ cells were spotted onto plates, overlaid with a nitrocellulose membrane, and grown overnight. Secreted CPY bound to the nitrocellulose membrane and could be subsequently detected by probing with anti-CPY antibodies (Fig. 4A). WT and *avl9*Δ showed no detectable CPY secretion, suggesting that loss of *AVL9* does not disrupt CPY sorting. As expected, *vps4*Δ cells missorted CPY; however, secretion of CPY was greatly diminished in *vps4*Δ *avl9*Δ cells. Quantification confirmed this difference, as *vps4*Δ *avl9*Δ showed significantly reduced CPY secretion compared to *vps4*Δ alone (Fig. 4B). Notably, CPY secretion was detectable in *vps4*Δ *avl9*Δ cells, but was not significantly different compared to WT or *avl9*Δ. Plate-based growth of WT, *avl9*Δ, *vps4*Δ, and *vps4*Δ *avl9*Δ showed no growth defects or temperature sensitivity, demonstrating that changes in CPY missorting are not due to differences in the number of cells (Fig. 4C). This finding represented the first instance of secretory disruption of by loss of *AVL9* that was not dependent on loss of *VPS1* or *APL2*.

**Figure 4:**
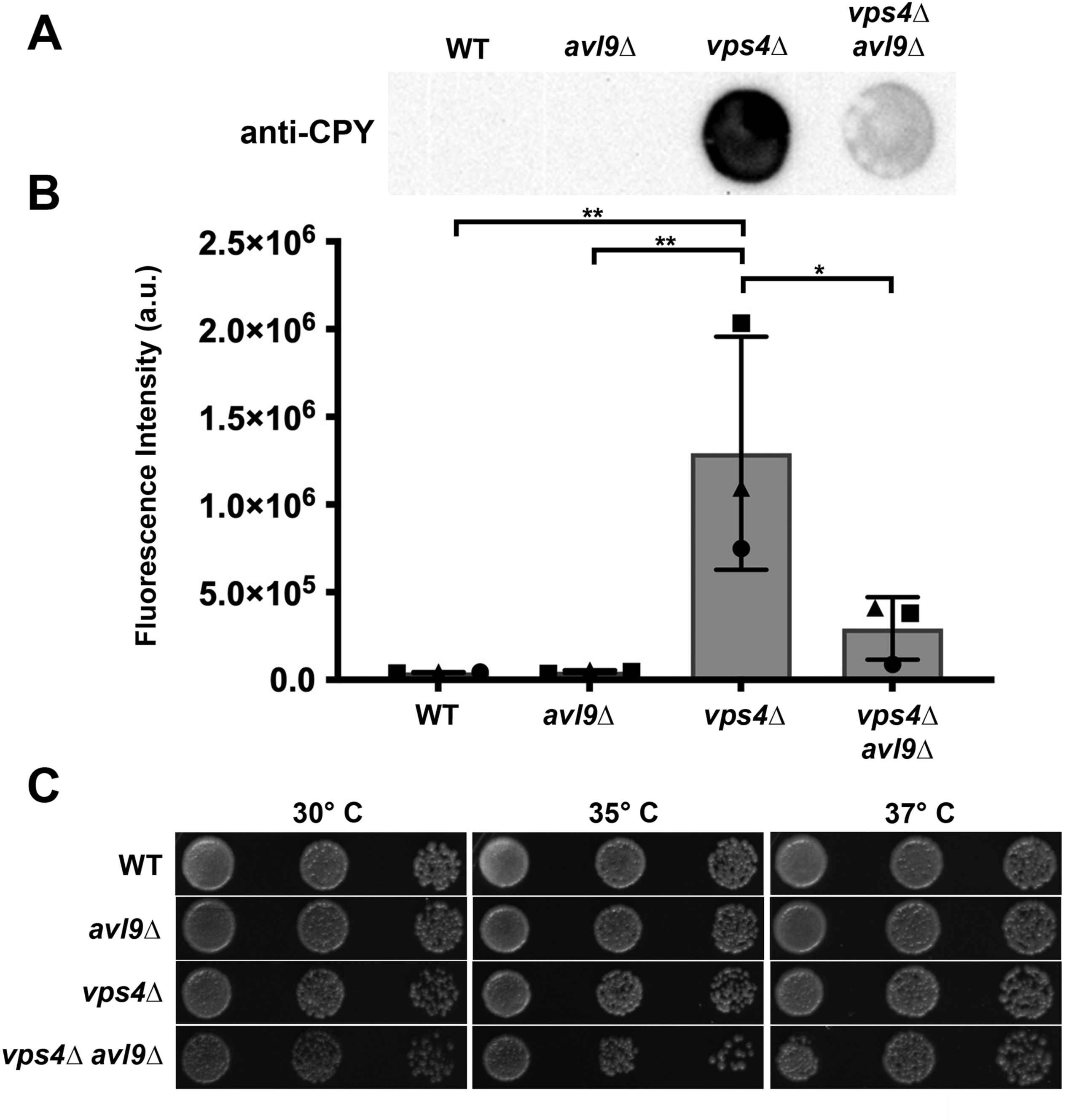
Carboxypeptidase Y (CPY) missorting in *avl9*Δ mutants. (A) WT, *avl9*Δ, *vps4*Δ, and *vps4*Δ *avl9*Δ cells grown to mid-logarithmic phase were concentrated to 2 OD_600_ in 50 μl, and 10 μl was spotted on to YPD plates. Culture spots were overlaid with a nitrocellulose membrane and grown for 18 h, and were then probed with anti-CPY. (B) Quantification of secreted CPY from cells shown in panel A. Values expressed as mean integrated density ± SD (in a.u.). One-way ANOVA with Tukey’s multiple comparisons test, n=3, (* p<0.05, ** p<0.01). (C) WT, *avl9*Δ, *vps4*Δ, and *avl9*Δ *vps4*Δ cells were grown to mid-logarithmic phase, serially diluted, and spotted onto YPD plates at the indicated temperature for 2 d.

A universal requirement for secretory vesicles is the presence of the v-SNARE Snc1 or its homologue Snc2, which drive vesicle fusion with the PM (Protopopov et al., 1993; Söllner et al., 1993). Following fusion with the PM, Snc1 is ubiquitinated and sorted onto endocytic vesicles, followed by recycling to the Golgi (Lewis et al., 2000). This well-documented process makes it an excellent marker for visually assessing secretion, endocytosis and recycling. We transformed WT and *avl9*Δ cells with a plasmid expressing GFP-Snc1 and examined localization of the cargo by fluorescence microscopy. WT cells displayed the expected distribution of GFP-Snc1, which polarized toward the bud. In contrast, *avl9*Δ cells showed internalized GFP-Snc1, with little fluorescence at the PM (Fig. 5A). To quantify this observation, we calculated the fraction of GFP-Snc1 at the PM and confirmed our visual assessment that *avl9*Δ significantly reduced the Snc1 surface-to-total ratio, confirming that loss of *AVL9* disrupted GFP-Snc1 recycling (Fig. 5B).

**Figure 5:**
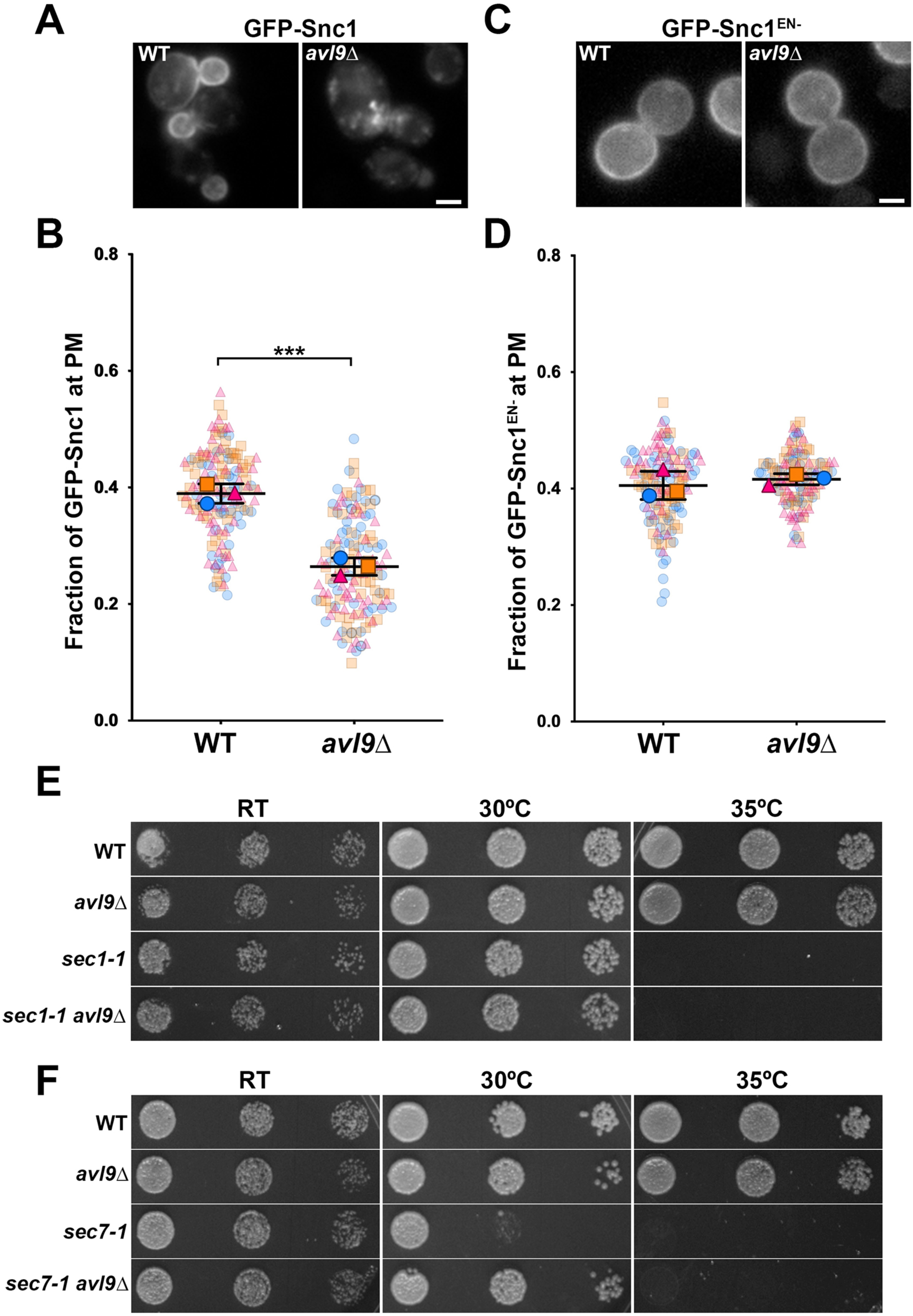
Effect of *avl9*Δ on recycling of Snc1. (A) WT and *avl9*Δ cells expressing GFP-Snc1 from a low-copy plasmid were imaged by fluorescence microscopy. (B) SuperPlot showing quantification of surface-to-total ratio of GFP-Snc1. Trial-level means (bars represent ± SD) are overlaid on scatter plots color-matched to trial-level data. Student’s t-test, n=3 biological replicates with 44-55 cells measured per condition for each trial (*** p <0.001). (C) WT and *avl9*Δ cells expressing GFP-Snc1^EN-^ from a plasmid were imaged using fluorescence microscopy. (D) Quantification of GFP-Snc1^EN-^ surface-to-total ratios as described in panel B. Student’s t-test, n=3 biological replicates with 39-52 cells measured per condition for each trial. (E,F) WT and *sec1-1* (E) or *sec7-1* (F) with and without *AVL9* were grown to mid-logarithmic phase, serially diluted, and spotted onto YPD plates at the indicated temperatures for 2 d.

While our results suggest that Avl9 contributes to Snc1 trafficking, these data alone do not distinguish between a role in secretion (TGN-to-PM) versus recycling from endosomes. To assess which pathway Avl9 acts on, we relied on an endocytosis-defective mutant of Snc1 (Snc1^EN-^). A V40A mutation in the sorting signal of Snc1 blocks its endocytosis, but not its exocytosis, resulting in GFP-Snc1^EN-^ becoming trapped at the PM and isolating its secretion from recycling (Grote et al., 1995; Lewis et al., 2000). When we expressed GFP-Snc1^EN-^ in WT and *avl9*Δ cells, we found that both strains showed an identical distribution of GFP-Snc1^EN-^, which was unpolarized at the PM and was observed on very few internal structures (Fig. 5C). Quantification of the surface-to-total fractions showed no significant differences in distribution of GFP-Snc1^EN-^ (Fig. 5D), suggesting that *AVL9* is required for recycling of post-endocytic cargos rather than secretion.

Given our earlier finding of negative genetic interactions between *AVL9* and *SEC4*, we next assessed whether the temperature sensitivity phenotype was exclusive to *SEC4* in the secretory pathway. We selected *SEC7* and *SEC1* because they are both essential genes that act at different stages of the secretory pathway. Sec7 is an Arf1 GEF critical for the activation of Arf1 and generation of vesicles at the TGN, and operates early in secretion(Franzusoff et al., 1991; Richardson et al., 2012). *SEC1* is a SNARE-binding protein that also interacts with the exocyst subunit Sec6 to help drive vesicle fusion at the PM during the final stages of secretion (Hashizume et al., 2009; Morgera et al., 2012). We deleted *AVL9* in temperature-sensitive mutants of both genes (*sec7-1* and *sec1-1*) (Novick and Schekman, 1979), and assessed growth phenotypes to determine whether *avl9*Δ altered temperature sensitivity as it did for *sec4-8* (Fig. 3A). We found that while *avl9*Δ did not alter *sec1-1* temperature sensitivity, it rescued *sec7-1* temperature sensitivity (Fig. 5E-F). Our finding of a positive genetic interaction between *avl9*Δ and *sec7-1* would suggest a degree of independence between these two genes, potentially functioning in separate routes to the PM that both require Sec1.

Our finding that *avl9*Δ disrupted Snc1 recycling prompted us to consider how Avl9 interacts with known recycling pathways. Previous studies showed that Snc1 is recycled through three independent routes mediated by the proteins Rcy1, Snx4, and Vps35 (Best et al., 2020). Rcy1 operates early in the endolysosomal pathway to recycle GFP-Snc1 and other cargos to the Golgi and PM, while Snx4 coordinates recycling from LE and Vps35 is a core subunit of the retromer complex involved in endosome-to-TGN retrieval (Best et al., 2020; Chen et al., 2005; Chen et al., 2011; Ma et al., 2017; MacDonald and Piper, 2017; Restrepo et al., 2007). We deleted *AVL9* in *rcy1*Δ, *snx4*Δ and *vps35*Δ strains and compared GFP-Snc1 localization and surface-to-total ratio in WT, single mutant, and double mutant cells. Both *avl9*Δ and *rcy1*Δ showed a similar loss of GFP-Snc1 at the PM with a corresponding increase in internal accumulation, while *rcy1*Δ *avl9*Δ showed a stronger defect than either single mutant; GFP-Snc1 fluorescence also appeared dimmer in the double mutant compared to either of the single gene deletions (Fig. 6A). The fraction of GFP-Snc1 at the PM similarly showed a significant decrease in *rcy1*Δ and *avl9*Δ compared to WT, and the *rcy1*Δ *avl9*Δ double mutant resulted in a further decrease in surface-to-total ratio that was significant compared to both single deletion mutants or to WT (Fig. 6B), suggesting that Avl9 and Rcy1 may operate in parallel pathways. The *snx4*Δ and *avl9*Δ cells showed a similar defect in GFP-Snc1 recycling, while the double *snx4*Δ *avl9*Δ mutant showed larger GFP-Snc1 structures that did not appear to be dimmer than internal structures in the single mutants (Fig. 6C). PM fractions for *snx4*Δ and *avl9*Δ showed similar, significant decreases compared to WT (Fig. 6D). The PM fraction of the *snx4*Δ *avl9*Δ double mutant was significantly decreased compared to *avl9*Δ, but not to *snx4*Δ, suggesting that since Snx4 operates late in the endolysosomal pathway, Snx4 may be more critical as a backup mechanism to recover GFP-Snc1 from LEs. (Ma et al., 2017). Lastly, we observed that *vps35Δ* showed accumulation of GFP-Snc1 on internal structures and at the PM, while *avl9*Δ or *vps35*Δ *avl9*Δ showed GFP-Snc1 on internal structures with reduced PM fluorescence (Fig. 6E). While both *vps35* and *avl9*Δ showed a decreased GFP-Snc1 PM fraction compared WT, the defect was not as severe in *vps35*Δ and we observed no difference between *vps35*Δ *avl9*Δ and either single deletion mutant (Fig. 6F). We reasoned that since Vps35 plays a relatively minor role in GFP-Snc1 recycling, and operates at a similar stage as does Snx4, this lack of difference may be due to Snx4 still functioning to recover downstream GFP-Snc1 from LEs (Best et al., 2020).

**Figure 6:**
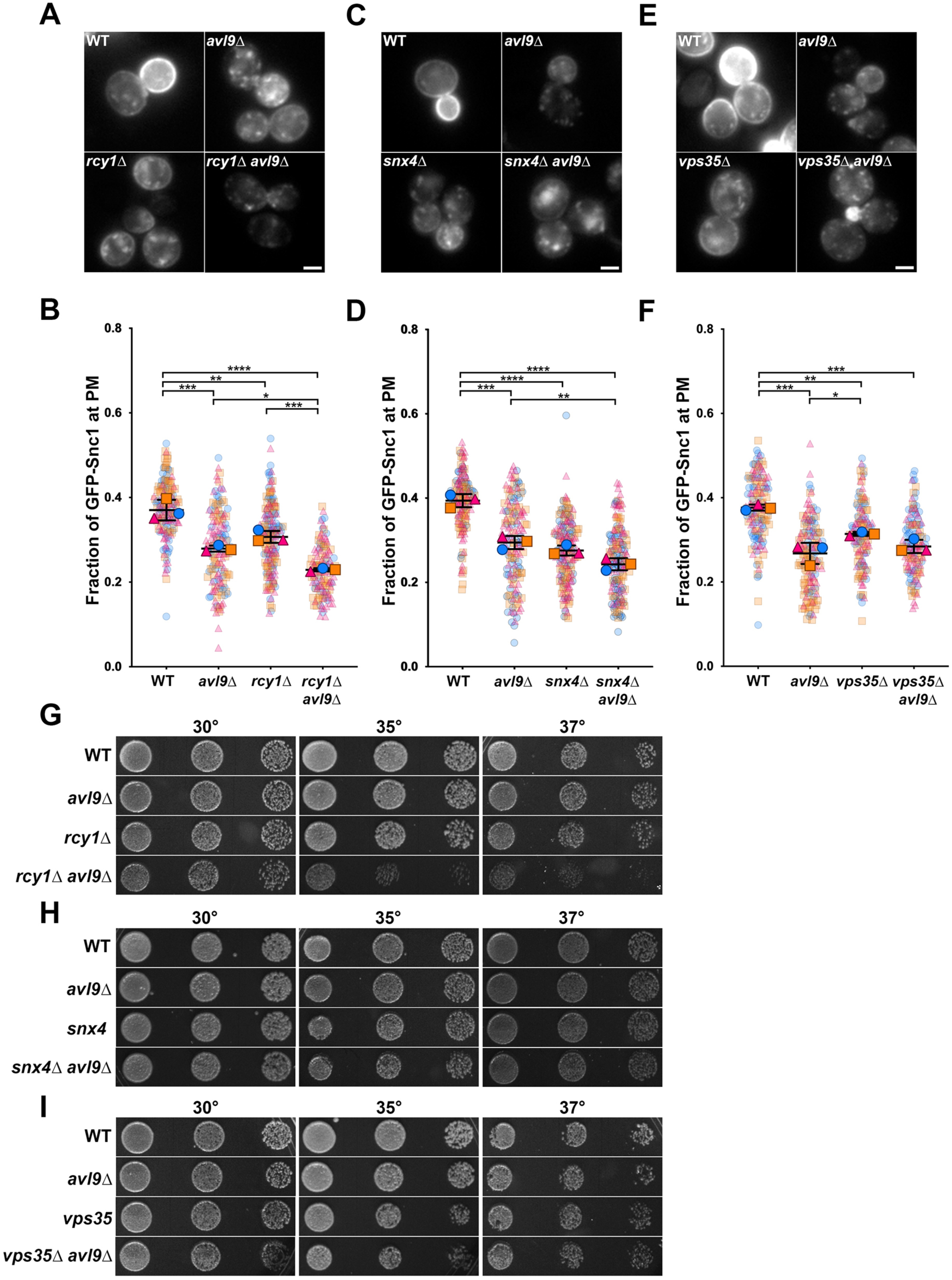
GFP-Snc1 in recycling mutants. (A) WT, *avl9*Δ, *rcy1*Δ and *rcy1*Δ *avl9*Δ cells expressing GFP-Snc1 from a low-copy plasmid were imaged by fluorescence microscopy. (B) SuperPlot showing quantification of surface-to-total ratio of GFP-Snc1. Trial-level means (bars represent ± SD) overlaid on scatter plots color-matched to trial level data of WT, *avl9*Δ, *rcy1*Δ and *rcy1*Δ *avl9*Δ cells. One-way ANOVA with Tukey’s multiple comparisons test, n=3 biological replicates with 35-64 cells measured per condition for each trial (* p<0.05, ** p<0.01, *** p<0.001, **** p<0.0001). (C) WT, *avl9*Δ, *snx4*Δ and *snx4*Δ *avl9*Δ cells expressing GFP-Snc1 from a plasmid were imaged using fluorescence microscopy. (D) SuperPlot as described in panel B showing quantification of surface-to-total ratio of GFP-Snc1 in WT, *avl9*Δ, *snx4*Δ, and *snx4*Δ *avl9*Δ. One-way ANOVA with Tukey’s multiple comparisons test, n=3 biological replicates with 40-70 cells measured per condition for each trial (** p<0.01, *** p<0.001, **** p<0.0001). (E) WT, *avl9*Δ, *vps35*Δ and *vps35*Δ *avl9*Δ cells expressing GFP-Snc1 from a plasmid were imaged using fluorescence microscopy. (F) SuperPlot as described in panel B showing quantification of surface-to-total ratio of GFP-Snc1 in WT, *avl9*Δ, *vps35*Δ, and *vps35*Δ *avl9*Δ. One-way ANOVA with Tukey’s multiple comparisons test, n=3 biological replicates with 35-66 cells measured per condition for each trial (* p<0.05, ** p<0.01, *** p<0.001). (G-I) WT and *rcy1*Δ (G), *snx4*Δ (H) or *vps35*Δ (I) cells with and without *AVL9* were grown to mid-logarithmic phase, serially diluted, and spotted onto YPD plates at the indicated temperatures for 2 d.

Lastly, we examined temperature-dependent growth of *rcy1*Δ, *snx4*Δ, and *vps35*Δ mutants combined with *avl9*Δ to determine whether loss of *AVL9* showed genetic interactions with any of the known recycling pathways. We observed that neither *avl9*Δ nor *rcy1*Δ alone displayed temperature sensitivity; however, *rcy1*Δ *avl9*Δ double mutant cells showed decreased growth compared to WT or either single mutant at 35°C and 37°C (Fig. 6G). This genetic interaction between *RCY1* and *AVL9* further suggests that Avl9 operates early in the endolysosomal pathway, and possibly in parallel to Rcy1. This finding is further supported by the lack of any observable temperature sensitivity in *snx4*Δ *avl9*Δ or *vps35*Δ *avl9*Δ compared to either single mutation or to WT (Fig. 6H-I).

## Discussion

Based on previous descriptions of Avl9 involvement in secretion, we hypothesized that *AVL9* should show genetic interactions and potential co-localization primarily with Sec4 as the major Rab GTPase involved in exocytosis. We were thus surprised at the breadth of co-localization between Avl9 and Rabs acting along different trafficking pathways, as well as genetic interactions between *AVL9* and both secretory and endosomal Rabs. The finding that not all Avl9 co-localized with Sec4 suggests that there is a degree of independence between these two proteins, and may indicate that they have independent roles with a strong degree of overlap. Previous studies demonstrated the existence of two classes of secretory vesicles in yeast, light and dense, with possibly different routing mechanisms (Harsay and Bretscher, 1995; Harsay and Schekman, 2002). Indeed, missorted CPY appears to be packaged into different classes of vesicles based on where in its sorting pathway the disruption occurs, with missorted CPY being packed into light vesicles in *vps1*Δ, dense vesicles in *vps10*Δ and into both light and dense vesicles in *vps4*Δ (Harsay and Schekman, 2002). How are these missorted vesicles, along with endosomally-derived recycling vesicles, converted to secretory vesicles that can be directed to and fuse with the PM? Presumably, all secretory vesicles should require Sec4 to interact with exocyst at the PM, regardless of whether they originate from the TGN or are directed to the PM from endosomes, and our data suggest that Avl9 may be involved in coordinating the conversion of endosomal-derived vesicles into exocytic vesicles.

Genetic interactions and spatial coincidence between Avl9 and the Rab5 isoforms, along with the closely-related Ypt10, were an unexpected observation. Vps21 is the primary Rab5 isoform, driving delivery of both endocytic vesicles and TGN-derived vesicles to the endosome, with the other isoforms showing redundant functions and situational regulation (Borchers et al., 2023; Langemeyer et al., 2020; Peplowska et al., 2007; Singer-Krüger et al., 1994). Ypt52 is negatively regulated in a poorly-understood manner by interactions with Roy1 and Skp1 in its GDP-bound form under physiological conditions(Liu et al., 2011). Normally, *vps21*Δ results in growth defects and disruption of vesicle trafficking within the endolysosomal pathway. However, deletion of *ROY1* in *vps21*Δ rescues these defects due to disinhibition of Ypt52. Ypt53, the isoform with the strongest similarity to Vps21, is dispensable and is downregulated at the translational level under normal physiological conditions. When cells are subjected to nutrient stress, *YPT53* expression is upregulated; Ypt53 performs the same function as Vps21 and can completely compensate for loss of *VPS21* (Nakatsukasa et al., 2014); Schmidt et al., 2017). The closely-related Ypt10, which contributes to transition of endosomes from early to late stages through interactions with the Ypt7 GEF Ccz1, contains a PEST motif that promotes proteasomal degradation and rapid turnover (Langemeyer et al., 2020; Louvet et al., 1999). Notably, we found that while *vps21*Δ showed a growth defect under normal conditions, *avl9*Δ did not worsen this phenotype; however, *vps21*Δ *avl9*Δ did show an increased sensitivity to monensin compared to *vps21*Δ alone (Fig. 3F). This phenotype is similar to that of the *vps21*Δ *ypt52*Δ mutant. Since Avl9 co-localized with Ypt52 but not with Vps21 or Ypt53, these findings suggest that Avl9 and Ypt52 may function together in a pathway that is independent of, but whose function is still partially redundant with, Vps21 as the major Rab5 isoform.

Our results suggest that *avl9*Δ results in defective sorting and recycling of endocytosed Snc1. While this could be the result of failed Snc1 delivery to the TGN as in other recycling mutants, our finding that *avl9*Δ alone does not affect secretion of Snc1 instead indicate a more direct involvement in recycling. Recent studies showed that endocytosed FM4-64 is delivered directly from the PM to a Sec7-positive TGN compartment that acts as an early endosome (Day et al., 2018). Later studies utilizing super-resolution microscopy suggested that this TGN/EE compartment is defined by the t-SNARE Tlg2 that partially overlaps with, but is separable from, the Sec7 compartment (Toshima et al., 2023). The Tlg2 compartment thus acts as the yeast sorting/recycling endosome. However, another study showed multiple routes of Snc1 recycling or retrograde trafficking mediated by Rcy1, Snx4 and retromer from distinct stages of the endolysosomal pathway that occurs prior to arrival at the TGN (Best et al., 2020). Importantly, several studies demonstrate recycling of cargo directly to the PM from both early and late endosomes, while bypassing the TGN (Laidlaw et al., 2022; MacDonald and Piper, 2017). Our findings largely align with these studies, suggesting an EE that participates in recycling and is separate from the TGN. Importantly, Snc1 recycling is perturbed in *avl9*Δ cells, while the endocytosis-defective Snc1^EN-^ mutant is still transported to the PM in *avl9*Δ cells, demonstrating that Avl9 acts upon recycling cargos after they have undergone endocytosis, and not upon cargos *en route* from the TGN to the PM. Moreover, we show that the Snc1 recycling defect seen in *avl9*Δ and *rcy1*Δ is enhanced in the double mutant. Additionally, *rcy1*Δ *avl9*Δ shows a temperature sensitivity phenotype not seen in *rcy*Δ or *avl9*Δ. We interpret the relationship between Rcy1 and Avl9 as being the likely result of Rcy1 and Avl9 characterizing independent recycling pathways to the PM from early endosomes. This model would also explain the positive genetic interactions seen between *avl9*Δ and *sec7-1* that seem at odds with the negative interactions described between *avl9*Δ and *sec4-8* or *sec1-1.* Both Sec4 and Sec1 work at late stages of vesicle secretion and are required for fusion of vesicles at the PM, In contrast, Sec7 is required for vesicle formation at the TGN. Thus, there might be competition for late secretory Rabs and fusion machinery between TGN-derived vesicles and Avl9-positive recycling vesicles. Loss of *AVL9* could eliminate this competition, allowing for greater access to the fusion machinery in cells with diminished capacity to generate essential secretory vesicles from the TGN.

Based on our findings, we propose a model wherein Avl9 directs vesicles originating from EEs toward the PM for secretion (Fig. 7). In this model, Ypt52- or Ypt10-positive vesicles bud from the EE, and Avl9 is recruited to these vesicles after they depart the EE. Next, Sec4 may arrive to these Avl9-positive recycling vesicles to convert them to secretory vesicles and render them competent for fusion with the PM. This exchange could operate as a mechanism of convergence for two trafficking pathways (direct endosome-to-PM recycling and TGN-to-PM secretion) that are generally regarded as independent. Similarly, co-localization of Avl9 with Ypt1 and Ypt31, which are both involved in recycling of GFP-Snc1, may be evidence of a similar means of converting vesicles of multiple origins into secretory vesicles (Chen et al., 2005; Chen et al., 2011; Sclafani et al., 2010). We present genetic and microscopic evidence for this model, and our future studies will examine temporal dynamics and biochemical interactions to better understand the precise sequence of events governing the role for Avl9 in endosomal recycling.

**Figure 7:**
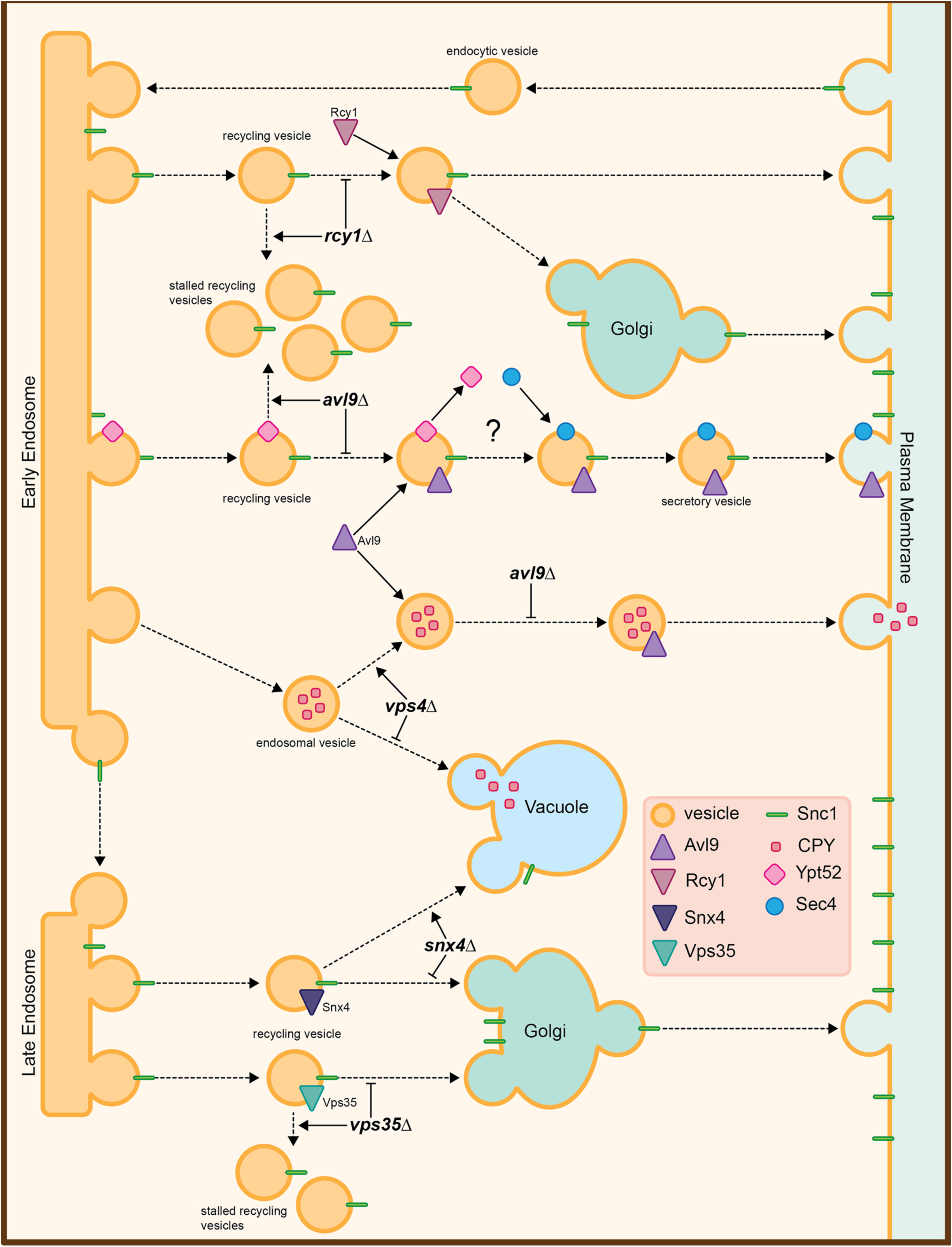
Model of proposed function for Avl9 in recycling and secretion. Snc1 on endocytic vesicles arrives at endosomes where it is sorted into recycling vesicles. After departing the endosome, these recycling vesicles acquire Sec4 and Avl9 before transport to the PM. Loss of *AVL9* blocks this progression, resulting in the accumulation of Snc1 on endosomal structures. CPY packaged into TGN/endosome-derived vesicles is normally transported to the vacuole. Loss of *VPS4* blocks fusion of these CPY-containing vesicles and sorts them into a secretory pathway in an Avl9-dependent manner.

The yeast endolysosomal network appears to be a carefully regulated system of convergent and divergent pathways. Cargos have several routes in and out of the endolysosomal pathway, and the full nature of how they are sorted and delivered to appropriate destination compartments remains an open question. Our study characterizes a novel role for Avl9 in mediating recycling and secretion of endosomal cargos. While additional studies are required to understand the function of Avl9, our study suggests that the physiological function of Avl9 is related to recycling rather than canonical secretion.

## Methods

### Yeast strains and growth conditions

Strains and plasmids used are listed in Supplemental Tables 1 and 2, respectively. All yeast strains were derived from BY4741 or BY4742(Brachmann et al., 1998). Cells were cultured in YPD medium (yeast extract, peptone, 2% dextrose) or YNB medium (yeast nitrogen base plus ammonium sulfate) lacking uracil or lysine for plasmid maintenance. Strains were grown at 30°C unless otherwise noted. Genomic tagging and knockout strains were generated by PCR-based methods as described (Goldstein and McCusker, 1999; Longtine et al., 1998). Transformations were performed using the lithium acetate method (Schiestl and Gietz, 1989). All reagents were from ThermoFisher or Sigma Aldrich unless otherwise indicated.

### Plasmids

GFP-Snc1 and GFP-Snc1^EN-^ plasmids were generously provided by Dr. Hugh Pelham (Lewis et al., 2000). GFP-Rab constructs used to generate plasmids for this study were gifts from Dr. Ruth Collins (Buvelot Frei et al., 2006). GFP-tagged Rabs were amplified by PCR from their original plasmids using M13 forward and reverse primers, and subcloned by homologous recombination into pRS317 digested with EcoRI and XmaI (New England Biolabs, Ipswich, MA, USA) (Sikorski and Boeke, 1991).

### Fluorescence imaging and analysis

Fluorescence microscopy was performed using a DMi8 inverted fluorescence microscope (Leica Microsystems, Wetzlar, Germany) equipped with a 100×, 1.47 numerical aperture (NA) Plan-Apochromat oil immersion lens, a Flash 4.0 v3 sCMOS camera (Hamamatsu, Shizuoka, Japan), an LED3 fluorescence illumination system, 488 nm and 561 nm lasers, a W-View Gemini image splitting optical device (Hamamatsu), compatible filter sets for fluorescence and DIC imaging, and LAS X v3.7.6.25997 software (Leica). Single-color fluorescence images were captured using acquisition parameters consistent between strains across all trials within an experiment. Two-color images were captured using the W-View Gemini beamsplitter to simultaneously capture both channels, using acquisition parameters consistent for strains within an experiment. Acquired images were imported into ImageJ/Fiji 2.16/1.54p, and 16-bit image files were adjusted to identical minimum and maximum intensity levels within an experiment. Images were exported as 8-bit TIFF files and cropped in Adobe Photoshop.

### Fluorescence quantification

To quantify the PM fraction of GFP-Snc1, DIC mode was used to select unbiased fields of cells representing all stages of the cell cycle and imaged using methods described above. 16-bit images were then imported into ImageJ, and background-subtracted. The freehand tool was used to draw a region of interest (ROI) around the outer perimeter of the PM, followed by a second ROI immediately inside the PM. This generated total fluorescence integrated density (total) and cytosolic fluorescence (cytosolic) integrated density values for each cell. The formula 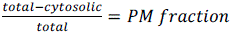 was then used to calculate to the fraction of GFP-Snc1 at the PM. All cells in each image were analyzed unless the cell touched the edge of the image, was dead or dying based on morphology in the DIC channel, or was out of focus(Prosser et al., 2016).

To quantify co-localization of Avl9-mCherry with GFP-Rabs, unbiased fields of cells were selected in DIC mode and both fluorescent channels were imaged simultaneously. Images were loaded into ImageJ as 16-bit image stacks, background subtracted, and unstacked. Using the green channel, the freehand tool was used to generate an ROI along the outer perimeter of each cell and the blow/lasso tool was used to generate an ROI of each puncta or subcellular structure observed. These ROIs were then opened in the corresponding red channel, where the number of total red punctae per cell (total punctae) along with the number of red punctae that coincided with a green ROI (co-localized punctae) were counted. The fraction of co-localized Avl9 punctae was then calculated as 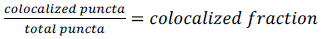.

### Plate-based growth assays

Genetic interactions were evaluated by spotting equivalent volumes of serially diluted strains grown to mid-logarithmic stage onto either YPD plates containing increasing concentrations of monensin or onto YPD plates that were incubated at increasing temperatures. Cells were inoculated in YPD medium and grown at 30°C overnight in a shaking incubator. Culture density was then measured at OD_600_ using a Genesys 10s Vis spectrophotometer (Thermo Fisher Scientific, Waltham, MA, USA). Cells were diluted to 0.4 OD_600_/ml and re-grown to mid-logarithmic phase (0.6-0.8 OD_600_/ml). Cells were then diluted to 0.25 OD_600_/ml, and a five-fold serial dilution series was performed (1:1, 1:5, 1:25). Cells were then transferred to 32-well pinning replicator plate (Dan-Kar Corp., Woburn, MA, USA) and a pronging device was used to transfer equivalent volumes of cells to appropriate plates in triplicate. YPD + monensin plates included drug concentrations of 0,12.5, 25, 50, 75 and 100 µM. Temperature sensitivity assays were performed at room temperature, 30°C, 35.5°C and 37°C as indicated. Plates were grown for 2-3 d and imaged using a FluorChem M gel documentation system (Bio-Techne, Minneapolis, MN, USA)

### Endpoint Assays

Endpoint assays were performed based on a protocol previously described in (Hung et al., 2018). Briefly, cells were grown overnight and density measured as described above for the plate-based assay. Cells were then diluted to a concentration of 0.0025 OD_600_/ml. A 2X stock of each monensin concentration (0, 6.5, 12.5, 25, 50, 100 µM) was prepared. In a 96-well dish, 100 µl of each 2X stock concentration of drug was added into two 6-well series for each strain (two technical replicates). 100 µl of cells were then added to the wells. The plate was placed in a SpectraMax i3x multi-mode plate reader controlled by SoftMax Pro v7.1.0 software (Molecular Devices, San Jose, CA, USA) and the initial concentration at OD_600_ was read. The plate was then transferred to a 30°C incubator for 18 h. The plate was removed from the incubator, gently vortexed to resuspend the cells, and the final OD_600_ density was read.

The raw data from both the initial and final reads for each experiment were exported to Microsoft Excel. Initial densities were subtracted from the final densities, and technical replicates were averaged. The growth at each concentration of monensin was then normalized to the control growth at 0 µM monensin.

### CPY missorting assay

Cells were cultured and grown to mid-logarithmic phase as described above for the plate-based assay. 2 OD_600_ of cells were then collected, washed with sterile YPD medium, and resuspended in 50 µl of media. 10 µl of cell suspension for each strain was then spotted onto a YPD plate by pipetting, and was allowed to dry at room temperature. A nitrocellulose membrane was then overlaid on the plate, and the plate was incubated at 30°C for 18 h. The nitrocellulose membrane was removed from the plate, rinsed with deionized H_2_O, and placed in blocking buffer [5% (w/v) skim milk powder in wash buffer (20 mM Tris-HCl pH 7.5, 150 mM NaCl, 0.1% Tween-20)]. The membrane was removed from the blocking buffer and probed with mouse anti-CPY [1:1000 dilution, monoclonal antibody 10A5-B5, ThermoFisher Cat. number A-6428] in blocking buffer for 1 h at RT. The membrane was then rinsed three times with wash buffer and probed with HRP-goat anti-mouse [1:5000, Jackson Laboratories Cat. Number 115-035-003] in blocking buffer for 1 h. The membrane was rinsed three times with wash buffer and developed for 2 min in an ECL solution. Chemiluminescence was then imaged with a FluorChem M gel documentation system. To quantify secreted CPY, the chemiluminescence images of membranes for each trial were imported into ImageJ as 16-bit files. Images were background subtracted, and the circle tool was used to draw an equal-sized ROI around each spot on the image for measurement of integrated density values.

### Statistical Analysis

All statistical analyses were performed using Prism 9 (GraphPad). Student’s t-test was used for GFP-Snc1 and GFP-Snc1^EN-^ in WT vs *avl9*Δ experiments. One- or two-way ANOVA with Tukey’s multiple comparisons test was used for Rab-Avl9 co-localization, Rab5Δ *avl9*Δ endpoint growth assay at 0 µM monensin, CPY missorting assay and GFP-Snc1 in *avl9*Δ recycling mutant experiments. Two-way ANOVA with Dunnet’s multiple comparisons test was used for the full Rab5Δ *avl9*Δ endpoint monensin growth assay across all monensin concentrations. All GFP-Snc1 quantitative microscopy experiments are displayed as SuperPlots in which trial level means were overlaid on individual trial points were color and shape coordinated to their trial level mean(Lord et al., 2020). Rab-Avl9 co-localization is displayed as SuperPlots of trial level means overlaid on violin plots of the individual trials.

## Acknowledgements

We would like to thank Drs. Hugh Pelham (Cambridge) and Ruth Collins (Cornell) for generous gifts of GFP-Snc1 and GFP-Rab plasmids used in this study. We are grateful to Dr. Jason Newton (VCU) and members of the Newton and Prosser labs for helpful comments and feedback.

---

**Supplemental Figure S1:**
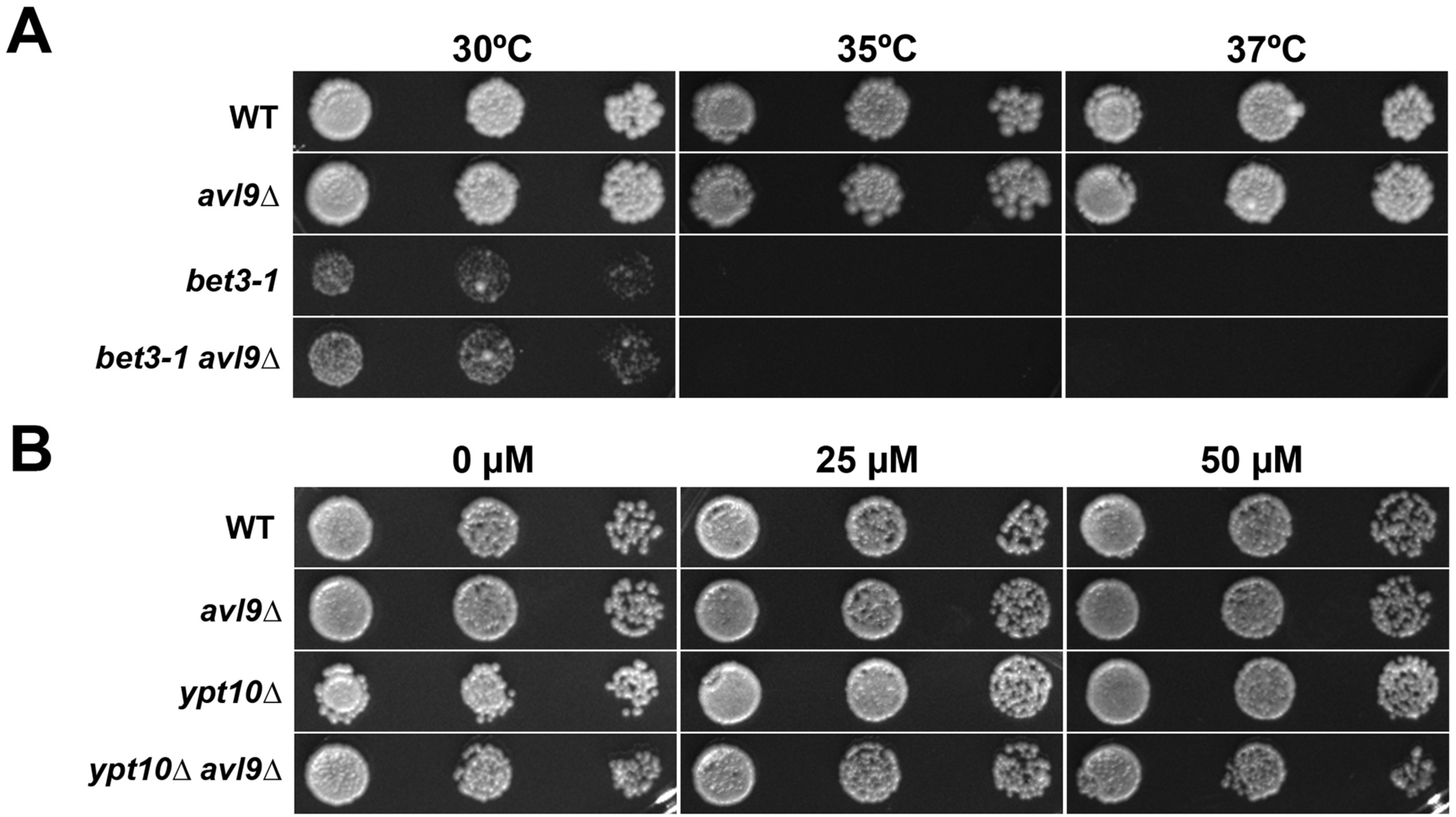
**Genetic interactions between *AVL9* and the TRAPP subunit *BET3* or the Rab5-like GTPase *YPT10*** (A) WT and *bet3-1* cells with and without *AVL9* were grown to mid-logarithmic phase, serially diluted, and spotted onto YPD plates at the indicated temperatures for 3 d. (B) WT and ypt10Δ with and without *AVL9* were grown to mid-logarithmic phase, spotted onto YPD plates containing 0, 25, and 50 μM monensin, and imaged after 3 d growth at 30°C.

**Supplementary Table S1:** Yeast strains used in this study.

| **Strain** | **Genotype** | **Source** |
| --- | --- | --- |
| DPY0046 | BY4741 *MATa his3Δ1 leu2Δ0 ura3Δ0 met15Δ0* | Brachmann et al., 1998 |
| DPY0047 | BY4742 *MATα his3Δ1 leu2Δ0 ura3Δ0 lys2Δ0* | Brachmann et al., 1998 |
| DPY1048 | BY4741 *vps21::KANMX6* | This study |
| DPY1050 | BY4741 *ypt52::KANMX6* | This study |
| DPY1052 | BY4741 *ypt53::KANMX6* | This study |
| DPY1053 | BY4742 *vps21::KANMX6 ypt52::KANMX6* | This study |
| DPY1058 | BY4742 *vps21::KANMX6 ypt53::KANMX6* | This study |
| DPY1059 | BY4741 *ypt52::KANMX6 ypt53::KANMX6* | This study |
| DPY1061 | BY4741 *vps21::KANMX6 ypt52::KANMX6 ypt53::KANMX6 lys2Δ0* | This study |
| DPY1847 | BY4741 *AVL9-GFP::HPHMX4 LEU2* | This study |
| DPY2395 | BY4741 *rcy1::KANMX6* | This study |
| DPY2396 | BY4741 *vps35::KANMX6* | This study |
| DPY2414 | BY4741 *avl9::NATMX6* | This study |
| DPY2416 | BY4742 *avl9::NATMX6* | This study |
| DPY2440 | BY4742 *ypt10::KANMX6* | This study |
| DPY2442 | BY4742 *ypt10::KANMX6 avl9::NATMX6 met15Δ0* | This study |
| DPY2486 | BY4741 *vps21::KANMX6 avl9::NATMX6 MET15* | This study |
| DPY2489 | BY4741 *ypt52::KANMX6 avl9::NATMX6 lys2Δ0* | This study |
| DPY2492 | BY4741 *ypt53::KANMX6 avl9::NATMX6* | This study |
| DPY2500 | BY4742 *sec4-8::KANMX6 lys2Δ0* | This study |
| DPY2502 | BY4742 *sec4-8::KANMX6 avl9::NATMX6 LYS2* | This study |
| DPY2522 | BY4741 *vps21::KANMX6 ypt52::KANMX6 avl9::NATMX6* | This study |
| DPY2523 | BY4741 *vps21::KANMX6 ypt53::KANMX6 avl9::NATMX6* | This study |
| DPY2524 | BY4741 *ypt52::KANMX6 ypt53::KANMX6 avl9::NATMX6 lys2Δ0 MET15* | This study |
| DPY2525 | BY4741 *vps21::KANMX6 ypt52::KANMX6 ypt53::KANMX6 avl9::NATMX6 lys2Δ0 MET15* | This study |
| DPY2541 | BY4741 *vps35::KANMX6 avl9::NATMX6* | This study |
| DPY2544 | BY4741 *rcy1::KANMX6 avl9::NATMX6* | This study |
| DPY2552 | BY4742 *bet3-1::KANMX6* | This study |
| DPY2566 | BY4742 *bet3-1::KANMX6 avl9Δ::NATMX6* | This study |
| DPY2843 | BY4741 *vps4::KANMX6* | This study |
| DPY2981 | BY4741 *AVL9-GFP::HPHMX4 LEU2 VPH1-mScarlet1::HIS3* | This study |
| DPY3073 | BY4741 *AVL9-GFP::HPHMX4 LEU2 IST1-mScarlet1::HIS3* | This study |
| DPY3075 | BY4741 *AVL9-GFP::HPHMX4 LEU2 VPS35-mScarlet1::HIS3* | This study |
| DPY3076 | BY4741 *AVL9-GFP::HPHMX4 LEU2 SEC7-mScarlet1::HIS3* | This study |
| DPY3077 | BY4741 *AVL9-GFP::HPHMX4 LEU2 SEC63-mScarlet1::HIS3* | This study |
| DPY3177 | BY4741 *snx4::KANMX6* | This study |
| DPY3190 | BY4741 *snx4::KANMX6 avl9::NATMX6* | This study |
| DPY3191 | BY4741 *vps4::KANMX6 avl9::NATMX6* | This study |
| DPY3603 | BY4742 *AVL9-mCherry::KANMX6* | This study |

**Supplementary Table S2:**
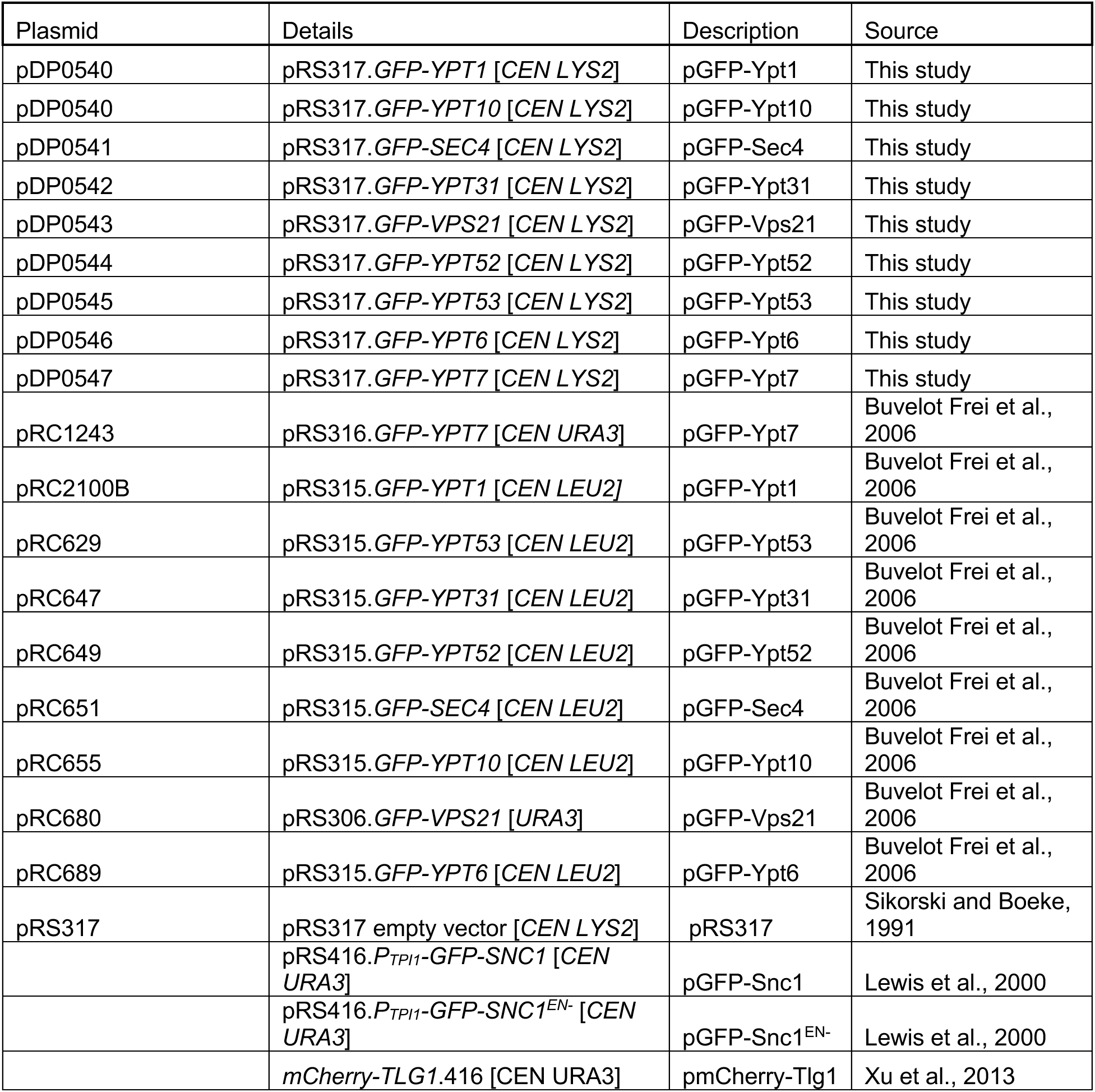
Plasmids used in this study.

